# “Galacturonic acid oxidation: a radical way to stick together”

**DOI:** 10.1101/2024.05.13.590697

**Authors:** Cyril Grandjean, Aline Voxeur, Salem Chabout, Stéphanie Boutet, Jérôme Pelloux, Sophie Bouton, Grégory Mouille

## Abstract

The middle lamellae (ML) of plant cells, enriched in Homogalacturonan (HG) is considered to function as a crucial ‘glue’ responsible for cell-cell cohesion (for review see ^1^). Mutants with defective HG content exhibit cell adhesion defects^2,3^. Despite advances, the mechanisms governing cell adhesion during plant development remain elusive. We previously hypothesized that cell-cell cohesion relies on cell wall integrity signaling, yet the specifics remain undefined^4^. OligoGalacturonans (OG), degradation products of HG, are prime candidates for ‘informing’ cells about ML status and thereby influencing cell adhesion. This integrity signal is crucial for adhesion homeostasis ^4^. OGs serve as signaling molecules, recognized by membrane-bound cell wall receptors. Notably, restoring adhesion in *qua2-1* mutants by modulating pectin response gene expression in *esmd/qua2* mutants underscores potential OGs’ importance^4–6^. Deciphering the diversity and role of endogenous OGs is imperative for understanding cell adhesion modulation. Our study aims to identify compounds in this signalling pathway that regulate cell adhesion. We focused on characterizing HG degradation products in dark-grown hypocotyls. Our findings highlight various oligomers, along with two key monomers: galacturonic acid and its oxidized form, galactaric acid. These monomers appear to play a pivotal role in controlling cell adhesion by indirectly enhancing the crosslinking of extensin, a cell wall structural protein. This crosslinking leads to the densification of extensin-based cell wall networks, ultimately restoring cell adhesion in defective mutants. Our research sheds light on the intricate interplay between HG degradation product monomer and cell adhesion mechanisms.

## Results and discussion

To explore the potential impact of endogenous HG-degradation products on cell adhesion we performed a qualitative and quantitative analysis of these oligosaccharides by UHPLC-HRMS (Ultra-High Performance Liquid Chromatography High Resolution Mass Spectrometry), as outlined in Voxeur et al. ^7^. Our focus was on etiolated hypocotyls from both the wild type Col0 and the adhesion mutant *qua2-1*. The endogenous HG fragments (extracted using 70% ethanol) are attributed to the activities of cell wall remodeling enzymes, including Pectin MethylEsterase (PME) and Polygalacturonase (PG). While both genotypes contained identical HG-degradation products, substantial quantitative differences were observed. Notably, *qua2-1* exhibited significantly reduced accumulation of each HG-degradation product compared to the wild-type (Figure 1), with oligomers content decreased by 29% to 50%, and galacturonic acid (GalA) and galactaric acid (GalAox) decreased by 84% and 30%, respectively. This was consistent with previous findings indicating a 50% reduction in homogalacturonans (HG) content in *qua2-1* compared to the WT.

**Figure 1.**
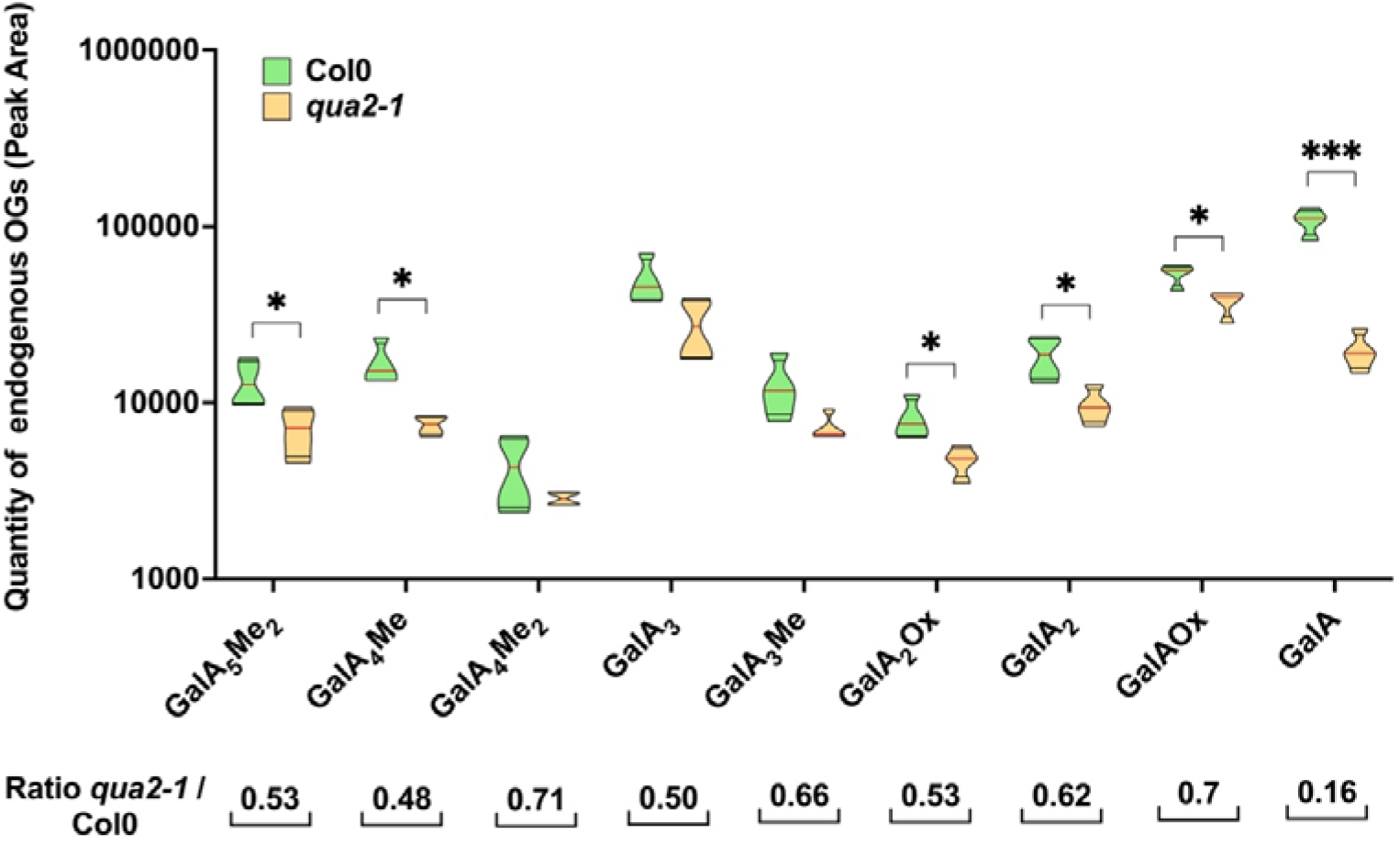
Quantity of endogenous OGs and monomers of *qua2-1* and Col0 dark-grown hypocotyls. The quantity of most of the OGs is lower in *qua2-1* compared to the wild type. The ratio of each OG of *qua2-1* compared to Col0 seems similar for most of the OGs (0.5 to 0.63 for the significant differences), except for the 2 monomers. Oligogalacturonides (OGs) are named as GalA_x_Me_y._ The number x in subscripts indicates the degree of polymerization, and y indicates the number of methylation or oxidation. GalA: galacturonic acid; Me: methyl ester group; Ox: oxidation. Red line represents the median, black line the quartiles (n = 4 biological replicates per genotype). *, *p* < 0.05, ***, *P* < 0.0001 Mann & Whitney test. Our analysis unveiled a previously unreported phenotype. In Col0, GalA was the dominant monomer, whereas in *qua2-1*, Oxidized Galacturonic acid (Galactaric) is the predominant monomer, as evidenced by the GalA/GalAox ratio. (1.99 for Col0 and 0.54 for *qua2-1*). To explore whether this difference might contribute to the distinct cell adhesion phenotypes of these genotypes, we cultivated dark-grown *qua2-1* hypocotyls in a medium supplemented with GalA. This treatment aimed to potentially restore the WT GalA/GalAox balance and assess its impact on *qua2-1* phenotypes.

The GalA-treated plants displayed a significant restoration of the cell adhesion defect, evident through ruthenium red staining (Figure 2A) and quantitative measurements of cell adhesion using the hypocotyls’ surface tortuosity score (Supplemental Figure 1A). GalA treatment restored adhesion in *qua2-1* mutants, yielding a score of -8.2, comparable to the WT score of 8.4.

**Figure 2:**
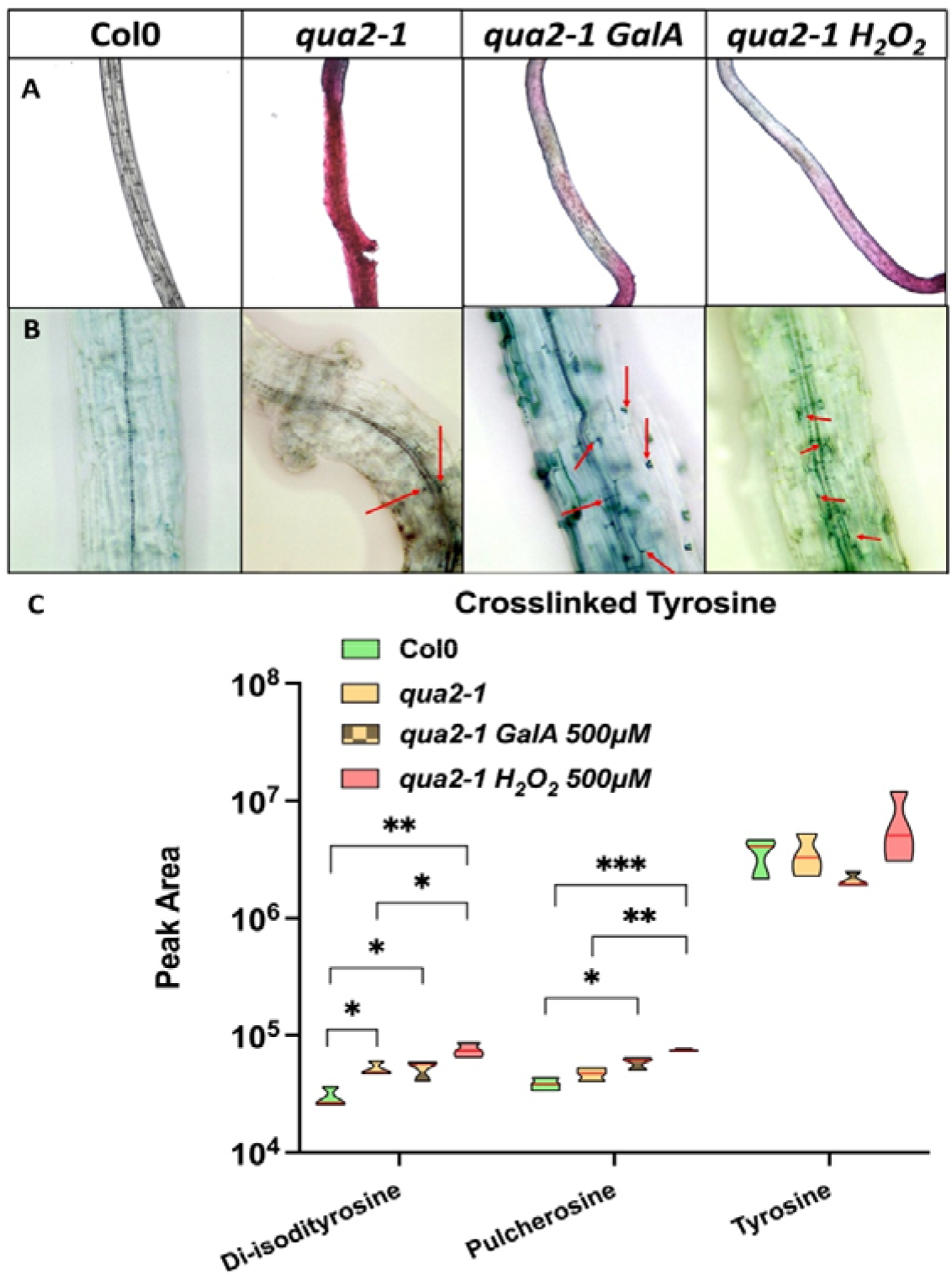
Galacturonic acid or hydrogen peroxide treatment restores cell adhesion defect of *qua2-1* mutant and increases the crosslinking ratio of tyrosine of cell wall proteins. A: Cell adhesion defect visualized by Ruthenium red staining. The GalA and H_2_O_2_ treatment restores the cell adhesion defect of *qua2-1*. B: Structural cell wall proteins crosslinked stained by Coomassie blue. (Red arrows pointed to hotspots of staining). The GalA and H_2_O_2_ treatment enhances the structural cell wall proteins crosslinked in *qua2-1*. C: Quantity of putative inter-molecular crosslinked tyrosine. The GalA treatment specifically increases the quantity of trimers in *qua2-1*. Red line represents the median (n = 3 biological replicates per genotype). * statistically significative difference, *p* < 0.05, **, *P* < 0.001, ***, *P* < 0.0001 T-Test.

In the context of investigating the impact of GalA-mediated restoration on the structure of the HG, we performed the analysis of the HG fine structure through enzymatic fingerprinting of the cell wall. This analysis revealed distinct patterns. Enzymatic fingerprinting of the cell wall using fungal endo-polygalacturonase (EPG) produced 93 HG fragments, including GalA monomer, with varying degrees of polymerization (DP) 1 to 13, and different methylation, acetylation, and oxidation signatures. Out of the fingerprinting results, we defined 5 classes, based on the nature of the substitution (Supplemental Figure 2). The wild-type exhibited a higher relative content of methylated HG fragments (49%) compared to *qua2-1* (37.7%), likely reflecting the defective pectin methyltransferase activity in *qua2-1*. This underscores the significance of HG fine structure, including methylation pattern, in mediating cell-to-cell adhesion. Intriguingly, GalA treatment induced minor changes in the relative content of *qua2-1* HG fragments (Supplemental Figure 2), suggesting that the quality of HG may not be the determinant for adhesion restoration. This highlights that cell adhesion recovery observed is independent of HG quality restoration in this case.

We next quantified the GalAOx in the control of the fingerprinting experiment, where the cell wall residue was incubated without the fingerprinting enzymes. It represents less than 1% of the total HG GalA content, explaining why it is not observed in the fingerprinting analysis. In this fraction, we identified cell wall localized monomers non-covalently bound to the cell wall.

Among these monomers, our results revealed a three-fold increase in Galacturonic Acid Oxidized (GalAOx) in *qua2-1* hypocotyls compared to Col0. Notably, when *qua2-1* hypocotyls were grown in a GalA-enriched medium, the amount of GalAOx was 30-fold higher than in control conditions (Supplemental Figure 3A). This shows that the exogenously added GalA is oxidized in the cell wall.

Considering these results, our next step was to analyze the consequences of applying GalAOx to *qua2-1* compared to iso-concentrations of GalA. Surprisingly, GalAOx did not restore the cell adhesion phenotype of *qua2-1* (Supplemental Figure 3B). This suggests that the reversion of the phenotype may be related to the conversion of GalA into GalAOx rather than GalAOx itself. As this conversion reaction is mediated by oxidases and generates H_2_O_2_, we further investigated the effects of applying H_2_O_2_ under conditions similar to those of GalA. H_2_O_2_-treated *qua2-1* seedlings were less stained by ruthenium red (Figure 2A), indicating that H_2_O_2_, like GalA, restores cell adhesion defects. To quantitatively assess the restoration by H_2_O_2_, we monitored the hypocotyls’ tortuosity levels (Supplemental Figure 1B). This analysis confirms that adhesion defects were reversed by H_2_O_2_ treatment, as shown by a tortuosity score of -8.1, which is not significantly different from that of Col0. To confirm that the reversion of cell adhesion defects in *qua2-1* was due to the direct effect of GalA and not a salvage pathway effect, we applied the same doses of glucuronic acid, the precursor of GalA, which did not affect cell adhesion (Supplemental Figure 1A). This demonstrates that both GalA and H_2_O_2_ treatments are closely linked to the mechanism suppressing cell adhesion defects in *qua2-1*.

Next, to understand how H_2_O_2_ mediates the restoration of cell adhesion, we hypothesized that it could be related to the crosslinking of Hydroxyproline-Rich Glycoproteins (HRGP), such as extensins, through their tyrosine residues, previously demonstrated in vitro ^9^. To test this hypothesis, we used Coomassie Blue staining to assess the extent of cross-linked proteins in the hypocotyls ^10, 11^. No staining was visible in Col0 hypocotyls (Figure 2 B), while staining was noticeable at the edge of some cells of *qua2-1*, where the cell adhesion defect is expected (Figure 2 B). This could be related to the higher relative abundance of GalAOx due to the higher conversion rate of GalA into GalAOx in *qua2-1* cell walls (Supplemental Figure 3A), which may lead to an increase in the amount of H_2_O_2_ and potentially increase crosslinking of proteins.

Seedlings grown in the presence of GalA or H_2_O_2_ (Figure 2B) revealed a much more intense and abundant labeling between the cells. Various types of intermolecular crosslinks of cell wall proteins have been described in previous studies ^9, 12-15^, including di-isodityrosine and pulcherosine (tetramer and trimer of tyrosine, respectively), which may yield crosslinked networks through C-O bonding catalyzed by peroxidases. Then in addition to the above-mentioned staining, we quantified the different forms of tyrosine -monomeric, trimeric (pulcherosine), and tetrameric (di-isodityrosine) – in the cell wall of dark-grown seedlings of wild-type and *qua2-1* grown in the presence or absence of GalA (Figure 2C). Di-isodityrosine levels were significantly higher in non-treated *qua2-1* mutant compared to the wild type, consistent with the increased staining observed with Coomassie blue. This suggests that in *qua2-1*, the mechanism of HRGP crosslinking is induced but not sufficient to maintain proper cell adhesion.

In *qua2-1*, the application of Galacturonic Acid (GalA) to dark-grown hypocotyls results in its conversion into Galacturonic Acid Oxidized (GalAox). This reaction likely involves a cell wall localized GalA oxidase from the berberine bridge enzyme family ^16^. This oxidation reaction liberates H_2_O_2._ Simultaneously, GalA treatment markedly increases the number of intermolecular extensin crosslinks, as indicated by the observed levels of di-isodityrosine and pulcherosine. These crosslinks are catalyzed by specific peroxidases ^17^. These peroxidases are presumed to utilize the available H_2_O_2_ generated from GalA oxidation along with extensin as electron donors, resulting in the formation of intermolecular bonds and subsequent restoration of the cell adhesion defect.

A prior study ^18^ documented the deposition of weakly or unesterified homogalacturonans (HGs) and extensins at the grafting interface in Arabidopsis, implicating the involvement of these two compounds in cell adhesion. This is in agreement with our results. We show that in the context of HG deficiency (*qua2-1)*, extensin crosslinking compensates for defective adhesion of the middle lamella. This study confirms the essential role of extensins as key mediators in cell adhesion.

## Experimental procedures

### Plant material and growth conditions

Col-0, *qua2-1 Arabidopsis thaliana* seedlings (150 seeds/genotypes) were grown in the dark at 21 °C on a solid medium Duchefa (1.1 g/L supplemented with CaNO_3_ 0.328 g/L, and 7.5 g/L of agar) at pH 5.7. This medium was only used for the endogenous OGs analysis. Otherwise, the *Arabidopsis thaliana seedlings* were grown in H_2_O, supplemented with 0.5 g/L of MES (Duchefa M1503.0100) buffered at pH 5.7 by adding KOH 0.04 M. For treated conditions, the liquid medium final concentration used was 500 μM of galacturonic acid, hydrogen peroxide, glucuronic acid, or galactaric acid. To synchronize the germination, seeds are cold-treated for 48h. Seeds were exposed to light for 4 h and then the plates were wrapped in 2 layers of aluminum foil and cultivated for 92h at 20°C. To not disturb the dark-grown condition, all cultures were harvested in a dark room under green light.

### Endogenous OG Content Analysis

An extraction was performed on fresh frost material of 150 dissected grown hypocotyls4-day-old dark-grown seedlings, with 70% ethanol, heated to 80 ° C during 1H. Endogenous OGs were analyzed by high-performance size exclusion chromatography (HP-SEC -MS-based) ^7^. Each endogenous OGs identified by the simple MS method was fragmented to confirm their identity. By this extraction and analysis, we identified 9 different OGs between a DP 1-5 decorated with different methylation or oxidation signatures.

### Ruthenium red staining

Fifty 4-day-old etiolated seedlings per genotype were stained with 0.5mg/ml ruthenium red solution for 2 min ^4^. After removing the solution, samples were washed twice with water. Images were acquired for each condition using an AXIOZOOM V16 ZEISS with an objective x 2.3.

### Tortuosity quantification -> Supp Data

Tortuosity is a property of a curve that is meandering, around the shape. We adapted a tool initially created to determine the cell shape dynamic by measuring the boundary curvature (Driscoll et al., 2012; Preetham Manjunatha, 2022). This tool was used to assess the tortuosity of the 2D binary shape of the hypocotyls and therefore relatively quantify the levels of cell detachment. Images of 4 or 5 hypocotyls were acquired for each condition using an AXIOZOOM V16 ZEISS with an objective x 2.3. The binary picture was obtained with Fiji software.

### Homogalacturonans fine structure analysis -> Supp Data

1.3U of commercial endo-polygalacturonase (*Aspergillus aculeatus*) was applied on 1-2 mg of alcohol insoluble residue (AIR) dry cell wall of 4-day-old dark-grown seedlings. A dry cell wall was obtained by washing the sample with ethanol 70 °C – 30 min at 80 °C and then with ethanol 96 °C – 30 min at 80 °C. Another washing step was performed twice with acetone for 20 min at room temperature. We performed digestion for 24 hours at 37 °C to hydrolyze the homogalacturonans fraction as much as possible. The fragments released were analyzed by high-performance size exclusion chromatography coupled with High-Resolution Mass Spectrometry (HP-SEC -MS-based) (Voxeur et al., 2019).

As a control for our experiment, we used the same material with only the inactivated enzyme and the buffer which is ammonium acetate 50 μM pH 5.

### Structural Cell wall proteins Crosslinks visualization

We used the protocol described by ^10,11^. Two biological replicates of about ten 4–day–old dark-grown seedlings were harvested and directly fixed in ethanol 96 % for 1 h at 80 °C and tranfered in 1% SDS solution at 80 °C for 24 hours to eliminate the proteins and keep only those that are crosslinked. The staining is then carried out with a solution of 0.1 % coomassie blue in H_2_O/ethanol/acetic acid (5/4/1) for 30 min and subsequently washed 3 times in H_2_O/ethanol/acetic acid (5/4/1) and makes it possible to visualize and localize the crosslinked proteins. Images of hypocotyls were acquired using an AXIOZOOM V16 ZEISS with an objective x 2.3.

### Cell wall cross-linked tyrosine identification and quantification

We sought to quantify trimeric, and tetrameric tyrosine from 4-day-old seedlings grown in the dark. A 6 N HCL hydrolysis was carried out on alcohol insoluble residue (AIR) and we analyzed the hydrolysate by HPLC HRMS (Ultimate 3000 Thermo QTOF IMPACTII Bruker). Chromatographic separation was carried out on a MACHEREY NAGEL Nucleoshell RP 18plus column (125 Å, 2.7 m, 2 mm X 100 mm), coupled to a Nucleoshell RP 18 plus guard column (125 Å, 2.7 m, 2 mm X 4 mm). The elution was carried out with a gradient of water with 0.1 % formic acid and acetonitrile with 0.1 % formic acid, with a flow rate of 400 μl / min and a column oven temperature of 40 °C.

The injection volume was set at 5 μl. ESI MS and MSMS detections are performed in positive and negative modes with the following ionization conditions: end plate offset voltage at 350 V, capillary voltage at 4500 V, nebulizer gas flow at 30 psi and drying time at 6 L / min for a source temperature of 250 °C. For the MS mode, the acquisitions are carried out over a mass range going from m / z 70 to m / z 800 mass unit with a scan speed of 1 Hz.

For ESI MSMS mode in DDA (Data-dependent acquisition) mode, a scan speed is set at 1 Hz to optimize the sensitivity and the mass range from m / z 70 am / z 800 mass units. The reference ions were deduced from the injection of tyrosine and dityrosine standards. For MSMS in DDA, the reference ions are 182.1076 (tyrosine), 361.1364 (di-tyrosine), 540.1971 (pulcherosine), 719.2550 (di-iso-di-tyrosine) over a collision energy range for fragmentation ranging from 20 to 100 eV depending on the masses selected.

HyStar 4.1 SR2 and DataAnalysis 4.4 software (Bruker Daltonics) were used for data acquisition, internal calibration from sodium formate cluster (Calibration by an HPC equation with a positive mass delta of 1.5ppm and a negative mass delta of 0.7ppm) and data visualization. Each identified molecule was fragmented by MS / MS and subjected to a match formula and fragmentation profile, searched with the SIRIUS Web module. We were able to identify 3 different forms of tyrosine ranging from the single amino acid to the tetramer Di-isodityrosine.

## Acknowledgments

The IJPB benefits from the support of Saclay Plant Sciences-SPS (ANR-17-EUR-0007). This work was supported by IJPB’s Plant Observatory technological platforms. Special thanks to Rawen Ben Malek for her critical reading.

## Authors contributions

G.M, S. Bouton and C.G. conceived the project and designed the experiments. C.G., A.V, S.C, S. Boutet performed the experiments. C.G. and G.M. wrote the manuscript. S. Bouton, A.V, J.P. contribute to the writing.

---

**Supplemental Figure 1:**
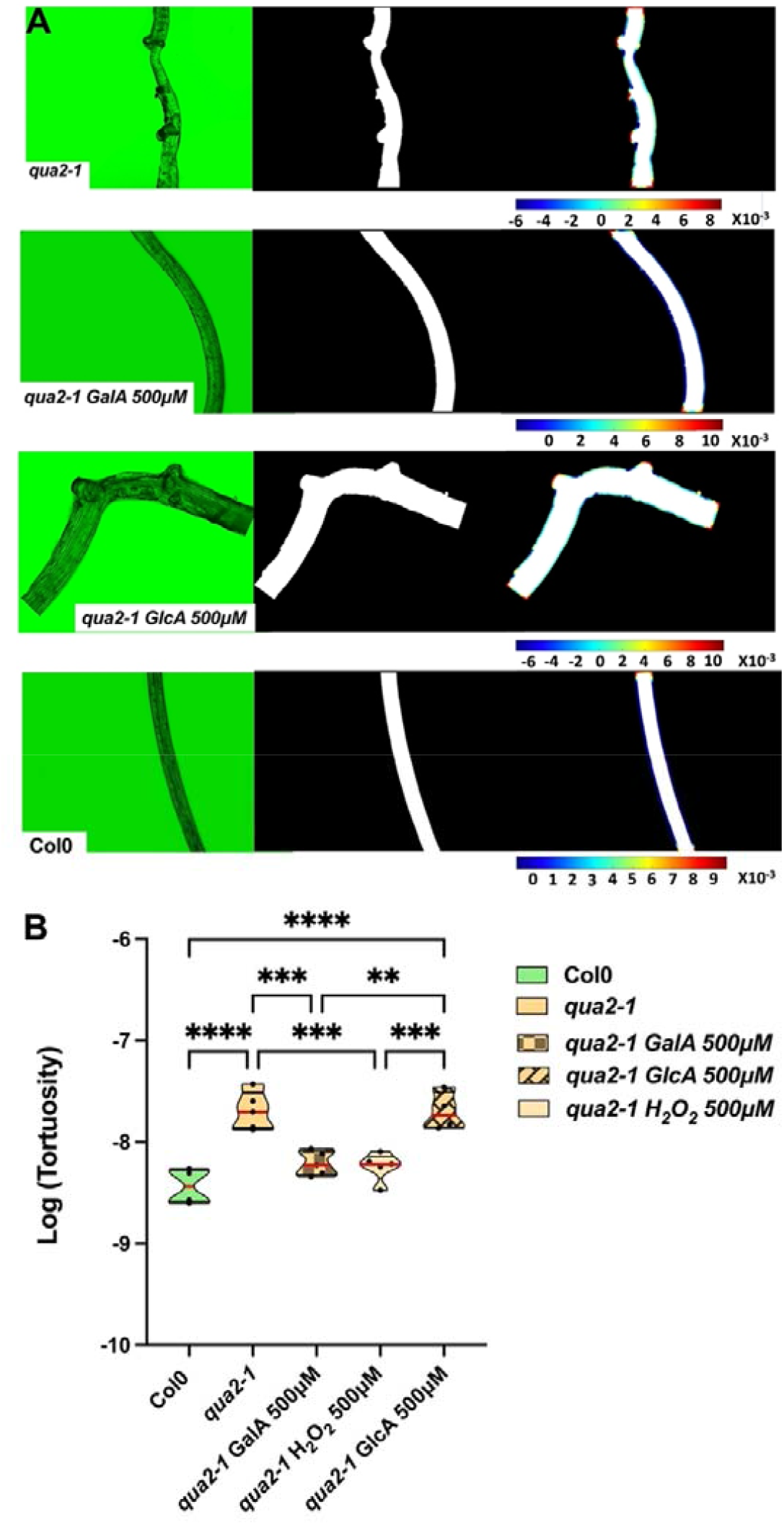
A new tool to quantify cell adhesion defect. (A) From left to right: Original picture (green channel only), binary image, analyze particle to get the contour of the shape the color scale corresponds to the level of curvature of the shape (Negative positive, or straight). The binary images are analyzed using code of MATLAB software according to (Driscoll et al., 2012; Preetham Manjunatha, 2021). (B) The tortuosity (corresponding to the shape curvature data/perimeter) of the 2D shape of *qua2-1 and qua2-1* treated with GlcA are significantly higher compared to the wild type. The GalA and H_2_O_2_ treatment on *qua2-1* significantly restored tortuosity. The treated *qua2-1* shows no significant difference compared to the wild type. In(A) **, P < 0.001, ***, P < 0.0001, ****, P < 0.00001, 1-way-ANOVA (Tukey’s multiple comparisons test, n ≥ 4 biological replicates per condition).

**Supplemental Figure 2:**
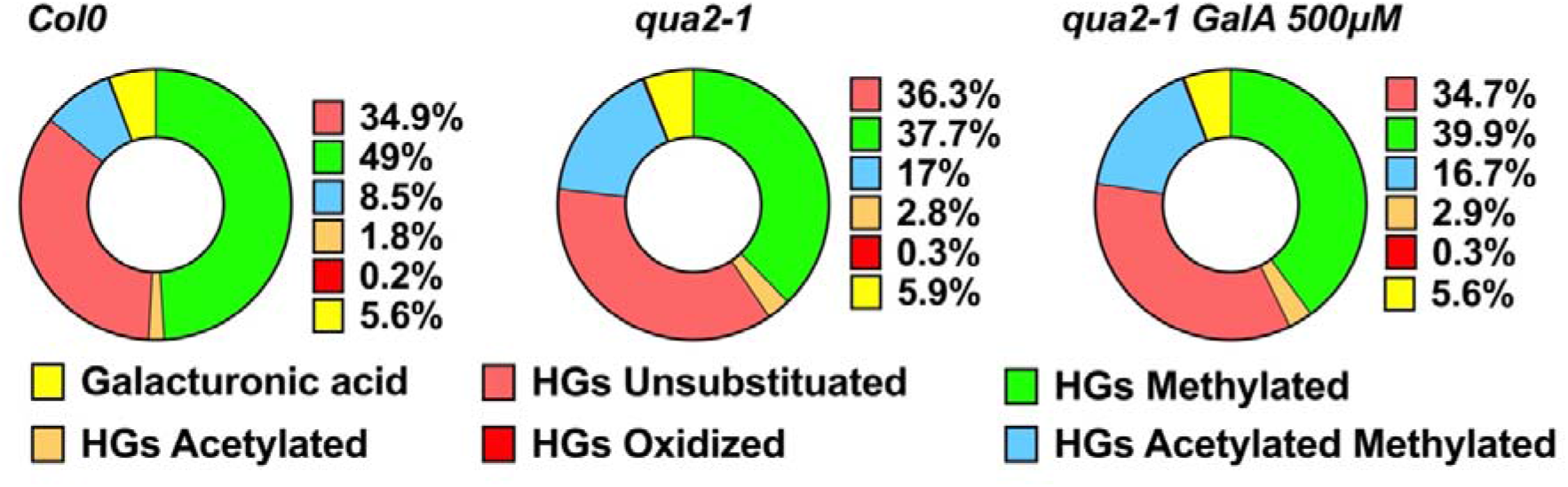
The effect of Galacturonic acid on the HG pattern of *qua2-1*. Relative content of homogalacturonans fragments family released by PG digestion cell walls *(n = 4 biological replicates per genotype)*. The GalA treatment induces only a slight change in *qua2-1* homogalacturonans pattern.

**Supplemental Figure 3:**
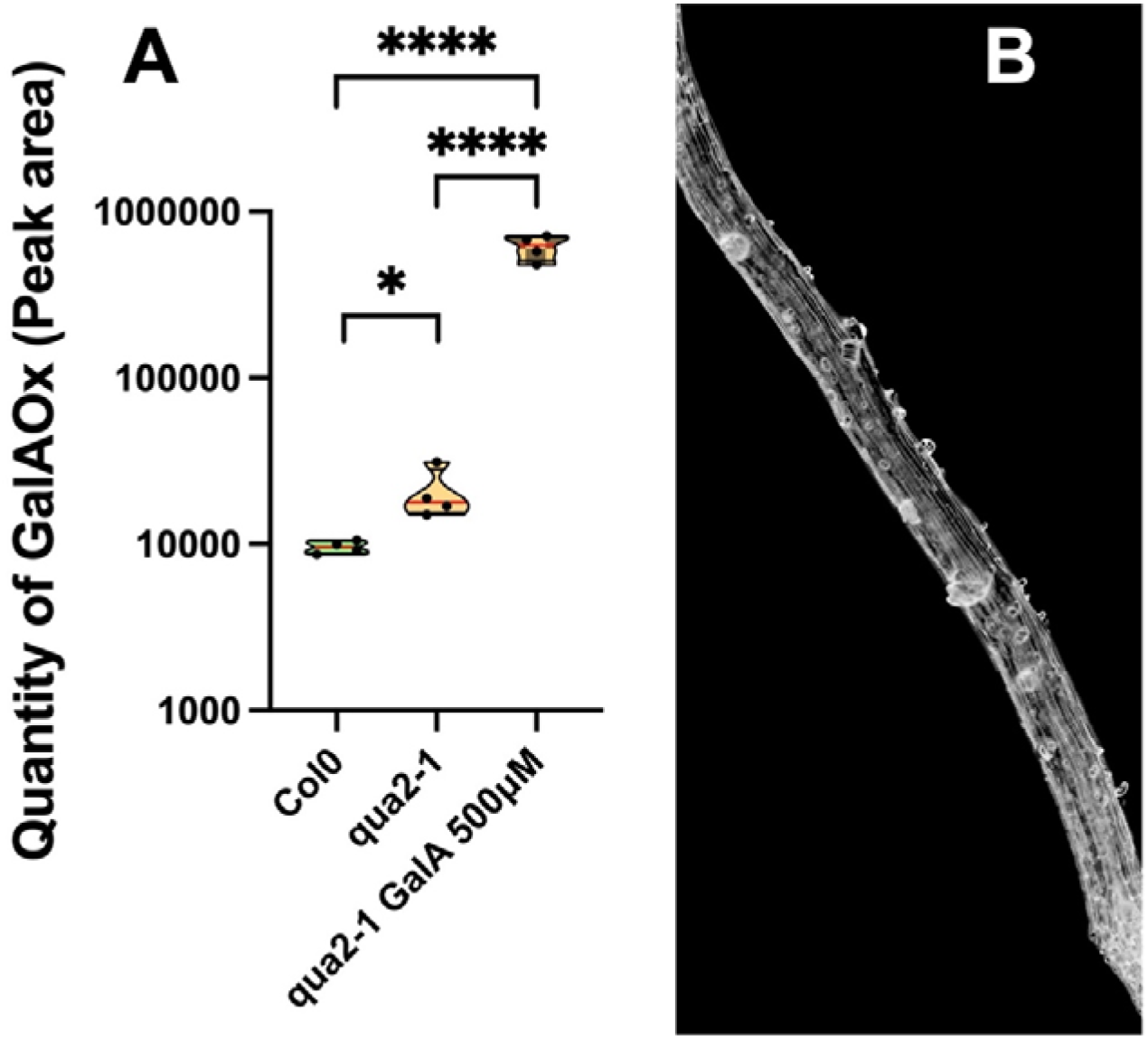
Galacturonic Acid is converted into galactaric acid. (A) Quantity of galactaric acid (GalAOx) trapped in the cell wall. The quantity of GalAOx is already higher in *qua2-1* compared to the wild type and multiplied by 30 after GalA treatment in *qua2-1*, suggesting that the added GalA is oxidized in the treated condition. Red line represents the median (n = 4 biological replicates per genotype). (B) Picture of 500μM GalAOx treatment on *qua2-1*. The GalAOx does not restore the cell adhesion suggesting is an intermediary of the mechanism we have highlighted. In (A) *, P < 0.05, ****, P < 0.00001, T-Test.

